# Glycopolymers stabilize protein folding and protein-protein interactions via enthalpic interactions

**DOI:** 10.1101/2025.08.11.669795

**Authors:** Sabrina M. Richter, Neal Brook, Alex J. Guseman

## Abstract

Macromolecular crowding is ubiquitous to physiological environments, perturbing the thermodynamics and kinetics of proteins via excluded volume and nonspecific chemical interactions. While crowding has been well-studied *in vitro* and in cells, the inert sugar polymers used to simulate crowding lack the chemical characteristics of biomolecules. Emerging studies guide the development of more relevant models of crowding in the cell, but little work has been done to discern crowding effects on proteins at the cell surface. Using ^19^F NMR, we measure how protein stability, folding, and intermolecular interactions are modulated by three glycopolymers abundant at the cellular exterior. Biologically relevant glycopolymers including heparin, hyaluronic acid, and mucin significantly stabilize folding of the N-terminal domain of the Drk-SH3 protein. These interactions are enthalpically stabilizing, emphasizing the importance of chemical interactions for biologically relevant crowders. We further show that these glycopolymers stabilize a homodimer formed by the A34F variant of GB1, demonstrating that biological crowders not only affect isolated proteins, but also influence how proteins interact with one another.

Crowding is more complex than simple ideas of volume exclusion suggest, and our work guides a more comprehensive understanding of protein crowding in the context of the glycocalyx, the last frontier of the cell.

## INTRODUCTION

Biological concentrations of macromolecules range from 100 – 400 g/L.^1^ In these crowded and heterogeneous environments, proteins are influenced by nonspecific interactions between macromolecules which are absent in dilute solutions^2,3^. These interactions, including hardcore steric repulsions and chemical interactions, modulate different aspects of protein functionality, including the equilibrium thermodynamics and kinetics of protein folding^4–8^, protein diffusion^9,10^, rates of catalysis^11–13^, and protein-protein interactions^14–17^. Hard-core repulsions manifest themselves entropically and result from excluded volume effects as two molecules are unable to occupy the same space.^2^ Hardcore repulsions stabilize the most compact state of the protein, which often is the folded state. Chemical interactions can be attractive or repulsive, depending on the chemical features of the macromolecules involved; repulsive chemical interactions are stabilizing to protein folding and protein-protein interactions, while attractive interactions are destabilizing.^3^ These interactions have been well studied *in vitro* and in living cells with a focus on how proteins fold and interact with each other in the cellular interior.^4,5,15,18–20^ Here, we look to expand this understanding to the last frontier of the cell, the glycocalyx.

The cellular glycocalyx is a dense mesh of glycopolymers that coats the outside of the cell, where it plays a fundamental role in processes including cell motility, adhesion, viral and bacterial infection, and oncogenesis.^21–23^ These functions are potentiated through its chemical and physical characteristics. The chemical composition of the glycocalyx is hallmarked by an abundance of proteoglycans and glycoproteins.^24^ Proteoglycans include a protein core attached to one or several linear glycosaminoglycan (GAG) polymers spanning hundreds to thousands of kDa and richly functionalized with negatively charged chemical groups.^25,26^ Glycoproteins consist of larger protein cores, to which smaller and often branched carbohydrates are anchored.^27^ Physically, the glycocalyx creates a barrier of entry to the cell membrane as it can extend tens of nanometers up to five micrometers beyond the cell surface.^23^ The glycocalyx varies in density, glycan composition, and thickness from organism to organism, and is also influenced by factors such as cell type, metabolism, and cell cycle state.^21–24^ Proteins that reside in the glycocalyx often undergo conformational exchange between an inactive state and an active state, and this interchange is tightly regulated.^28,29^

The structural and chemical complexity of the glycocalyx render it challenging to reconstitute, and studies which have aimed to simulate crowding effects at the cell surface have used the same synthetic sugar polymers routinely used to mimic crowding of the cellular interior.^30,31^ Here, we characterize the crowding effects of two GAG polymers, heparin and hyaluronic acid, and the highly glycosylated protein mucin, which together are representative of the heterogenous sugar polymers that occupy the exterior of the cell. Heparin is a 3-30 kDa polysaccharide comprised of repeating GlcUAα1-4GlcNAc disaccharide units, notable for its high levels of sulfonation.^32^ It is one of the most abundant constituents of the glycocalyx.^24^ Hyaluronic acid is a linear, carboxylated GAG made of repeats of GlcUA*β*1-3GlcNAc*β*1-4, distinguished for its exceptional length upwards of 5MDa.^26^ Mucins are a class of glycoproteins rich in serine and threonine residues which serve as the backbone for branched O-linked glycans.^27^ Here, we use ~15 kDa heparin, 50 kDa hyaluronic acid, and a heterogenous mixture of sialylated mucins isolated from porcine stomach.

Using two model systems, we compare the crowding effects of these charged glycopolymers relative to ficoll, a synthetic, uncharged sucrose polymer routinely used to generate crowded conditions. First, we measure how these crowders change the equilibrium thermodynamics and kinetics of protein folding, using the N-terminal SH3 domain of the *Drosophila* adapter protein Drk (SH3) as a model. SH3 exists in dynamic equilibria between a folded and unfolded state, and exchange between these states can be efficiently quantified by ^19^F NMR.^4,5,33,34^ To further characterize these physiological crowders, we measure their effects on the stability of a simple homodimer of the A34F variant of GB1.^35^ GB1 is a thoroughly studied globular protein, and dimerization of the A34F mutant is highly sensitive to changes in environmental conditions.^15,16,18,36,37^ The two-state equilibria of both systems make them amenable for NMR-based studies of protein dynamics.

## RESULTS

### Glycopolymers enthalpically stabilize the folded structure of SH3

SH3 has a standard state free energy of unfolding 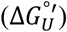 of ~0.5 kcal/mol.^5^ It coexists in a folded and unfolded state, and exchange between these states is slow on the NMR timescales.^38^ By fluorine-labeling the sole tryptophan residue of SH3 with 5-fluoroindole, populations of each state can simultaneously be quantified by ^9^F NMR (Fig **1a**). Using this two-state equilibrium, we measured how SH3 stability is perturbed by glycopolymers in concentrations as high as 300 g/L. Protein stability can be quantified by the modified standard free energy of unfolding (Eq. 1),

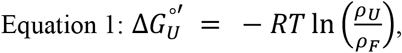

where R is the ideal gas constant and T is the absolute temperature. All the crowding conditions we generated stabilize the folded structure of SH3, and these effects are more prominent in physiological conditions compared with ficoll (Fig. **1b & 1c**, Fig. **S1** and Table **S1**). At 300 g/L, ficoll stabilizes SH3 by 0.31 ± 0.08 kcal/mol. At the same concentration, heparin and hyaluronic acid stabilize SH3 by 0.92 ± 0.08 kcal/mol and 1.5 ± 0.1 kcal/mol, respectively. Mucin also stabilizes protein folding, but concentrations above 200g/L could not be measured due to its extremely high viscosity. All three glycopolymers bias SH3 almost entirely to its folded state (Fig **1b** and **S1**).

**Fig 1.**
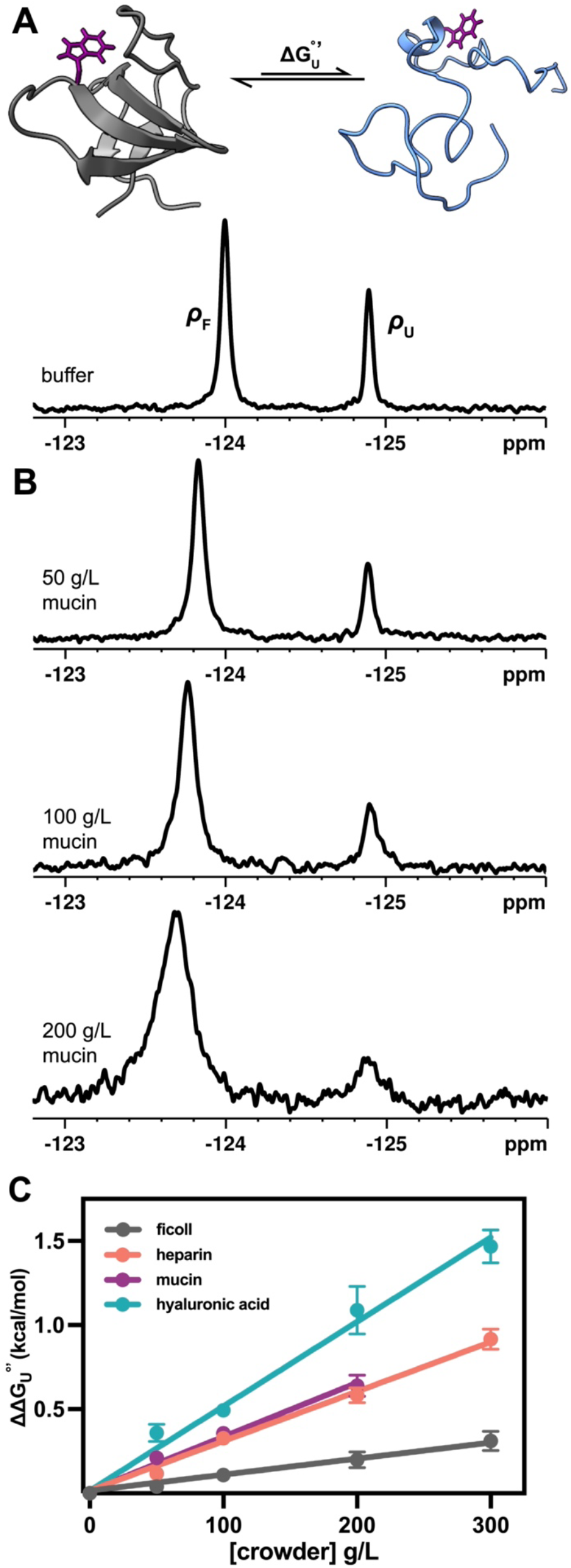
**a)** top, structure of SH3 (PDBID 2A36) in its folded (gray) and unfolded (blue) state, with fluorine-labelled W36 colored purple; bottom, ^19^F NMR spectra of SH3 in pH 7.0 buffer at 298K, with two resonances corresponding to folded and unfolded populations. **b)** representative stacked ^19^F NMR spectra of SH3 in increasing concentrations of mucin in pH 7.0 buffer, 298K. **c)** comparison of change in SH3 modified standard state free energies of unfolding relative to buffer across increasing crowder concentrations.

The resonances of SH3 broaden in crowded environments, and resonance broadening scales with crowder concentration (Fig. **S1, S2**, and Table **S2**). Nonspecific chemical interactions between proteins and macromolecules can manifest themselves via peak broadening, slower tumbling times and faster relaxation dynamics.^5,39^ To compare relaxation dynamics in different crowding environments, we estimated transverse relaxation times from linewidths.^40^ Estimated T2 times are ~2-4 fold faster in 300 g/L crowder conditions compared with buffer, and these effects are more outstanding in biological crowders compared to ficoll. Thus, we suspected that stabilization of SH3 by biological crowders arises substantially via chemical interactions.

To extract the entropic and enthalpic contributions of each crowding condition, we then measured the temperature dependence of 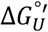 Data were fit to the integrated Gibbs-Helmholtz equation (Eq. 2) as described by Smith and colleagues^5^,

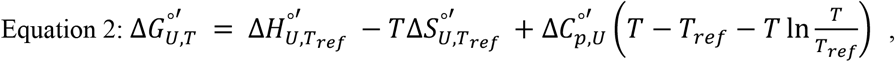

where *T*_*ref*_ refers to the melting temperature (when ρu = ρf). At 100 g/L, ficoll marginally stabilizes SH3 by 0.07 ± 0.06 kcal/mol, but both the entropic and enthalpic effects of ficoll are indistinguishable from buffer (Fig. **2** and Table **1**). Contrary to the idea of volume exclusion, we found that heparin, hyaluronic acid, and mucin entropically de*stabilize* SH3. Entropic destabilization of SH3 has previously been reported in other sugar polymers,^5,19,41^ and it is possible that the expected steric penalty of unfolding is not prevalent in crowder concentrations of 100 g/L used here. Enthalpically, the folded state of SH3 is favored in all three glycopolymers, and these effects are of greater magnitude than the entropic penalty of folding, leading to net stabilization. Chemical interactions thus have the dominant effect on protein stability.

**Fig 2.**
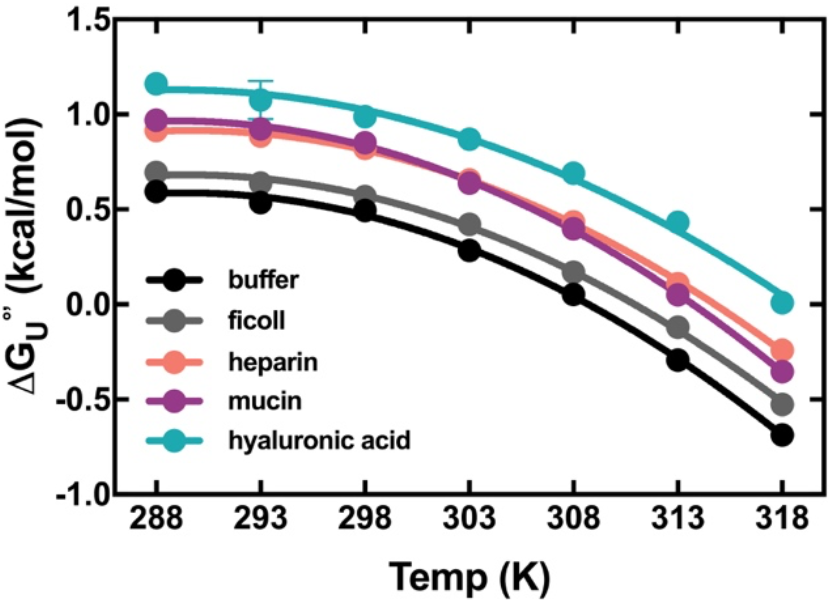
Temperature curves of SH3 in different crowding conditions. Data are shown as the mean ± SD of three measurements. Some error bars are not visible where the error is smaller than the data points.

**Table 1.**
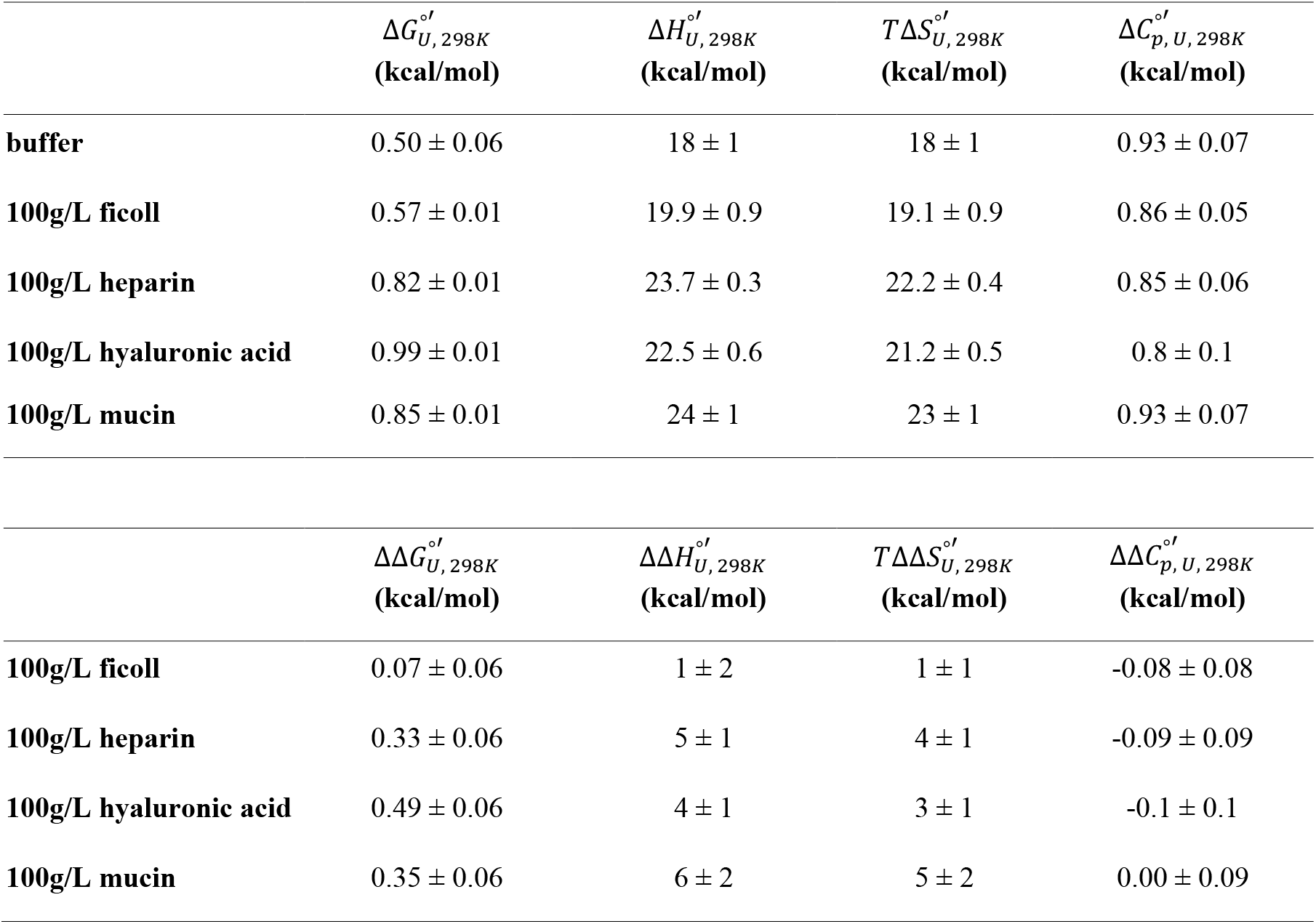
Thermodynamic parameters of SH3 at 298K.

| | $\Delta G_{U, 298K}^{o'}$<br>(kcal/mol) | $\Delta H_{U, 298K}^{o'}$<br>(kcal/mol) | $T\Delta S_{U, 298K}^{o'}$<br>(kcal/mol) | $\Delta C_{p, U, 298K}^{o'}$<br>(kcal/mol) |
| --- | --- | --- | --- | --- |
| **buffer** | $0.50 \pm 0.06$ | $18 \pm 1$ | $18 \pm 1$ | $0.93 \pm 0.07$ |
| **100g/L ficoll** | $0.57 \pm 0.01$ | $19.9 \pm 0.9$ | $19.1 \pm 0.9$ | $0.86 \pm 0.05$ |
| **100g/L heparin** | $0.82 \pm 0.01$ | $23.7 \pm 0.3$ | $22.2 \pm 0.4$ | $0.85 \pm 0.06$ |
| **100g/L hyaluronic acid** | $0.99 \pm 0.01$ | $22.5 \pm 0.6$ | $21.2 \pm 0.5$ | $0.8 \pm 0.1$ |
| **100g/L mucin** | $0.85 \pm 0.01$ | $24 \pm 1$ | $23 \pm 1$ | $0.93 \pm 0.07$ |
| | $\Delta\Delta G_{U, 298K}^{o'}$<br>(kcal/mol) | $\Delta\Delta H_{U, 298K}^{o'}$<br>(kcal/mol) | $T\Delta\Delta S_{U, 298K}^{o'}$<br>(kcal/mol) | $\Delta\Delta C_{p, U, 298K}^{o'}$<br>(kcal/mol) |
| **100g/L ficoll** | $0.07 \pm 0.06$ | $1 \pm 2$ | $1 \pm 1$ | $-0.08 \pm 0.08$ |
| **100g/L heparin** | $0.33 \pm 0.06$ | $5 \pm 1$ | $4 \pm 1$ | $-0.09 \pm 0.09$ |
| **100g/L hyaluronic acid** | $0.49 \pm 0.06$ | $4 \pm 1$ | $3 \pm 1$ | $-0.1 \pm 0.1$ |
| **100g/L mucin** | $0.35 \pm 0.06$ | $6 \pm 2$ | $5 \pm 2$ | $0.00 \pm 0.09$ |

### Glycopolymers have disparate effects on protein folding kinetics

Using 2D ^19^F EXSY experiments, we measured the folding (*k*f) and unfolding (*k*u) rates of SH3. Data were analyzed according to Farrow et al.^42,43^ In buffer, the folding rate of SH3 is 1.45 ± 0.07 s^−1^ and the unfolding rate is 0.65 ± 0.04 s^−1^ (Table **2**). The uncharged sugar polymer ficoll slows both processes with a folding rate of 1.1 ± 0.2 s^−1^ and unfolding rate of 0.43 ± 0.08 s^−1^. Extending this analysis to the glycopolymers, we found that heparin also slows both folding (0.99 ± 0.09 s^−1^) and unfolding (0.2 ± 0.1 s^−1^). For mucins and hyaluronic acid, the rate of folding increases to 1.75 ± 0.02 s^−1^ and 1.7 ± 0.3 s^−1^, respectively, while the rate of unfolding decreases to 0.35 ± 0.02 s^−1^ and 0.22 ± 0.05 s^−1^, respectively.

**Table 2.** Folding and unfolding rates of SH3 at 298K.

| | $k_f (\text{s}^{-1})$ | $k_u (\text{s}^{-1})$ |
| --- | --- | --- |
| **buffer** | $1.45 \pm 0.07$ | $0.65 \pm 0.04$ |
| **100g/L ficoll** | $1.1 \pm 0.2$ | $0.43 \pm 0.08$ |
| **100g/L heparin** | $0.99 \pm 0.09$ | $0.2 \pm 0.1$ |
| **100g/L mucin** | $1.75 \pm 0.02$ | $0.35 \pm 0.02$ |
| **100g/L hyaluronic acid** | $1.7 \pm 0.3$ | $0.22 \pm 0.05$ |

### Biologically relevant glycopolymers stabilize protein-protein interactions

Finally, we investigated how these biologically relevant glycopolymers bias protein-protein interactions using the T2Q A34F variant of GB1 (A34F), which exists in equilibria between a monomer and dimer.^35^ Y33 resides at the dimerization interface and experiences different chemical environments as a monomer and dimer; exchange between these states is slow, and fluorine-labelling the tyrosine residues of A34F yields two resonances for Y33 by ^19^F NMR, one corresponding to each state (Fig **3a & 3b**). We measured the stability of the A34F dimer in each crowding environment by using ^19^F NMR titrations at several protein concentrations (PT) between 15-500 µM. Peak integrations were used to determine the fraction monomer (*f*M) and dimer (*f*D) at each concentration, and dissociation constants (KD→M) were determined by fitting data to equation 3 in Prism using nonlinear least squares fit.

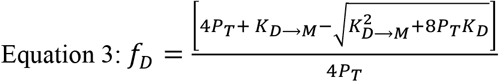

Binding energies could then be determined using equation 4,

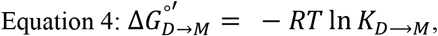

where R is the ideal gas constant and T is the absolute temperature.

**Fig 4.**
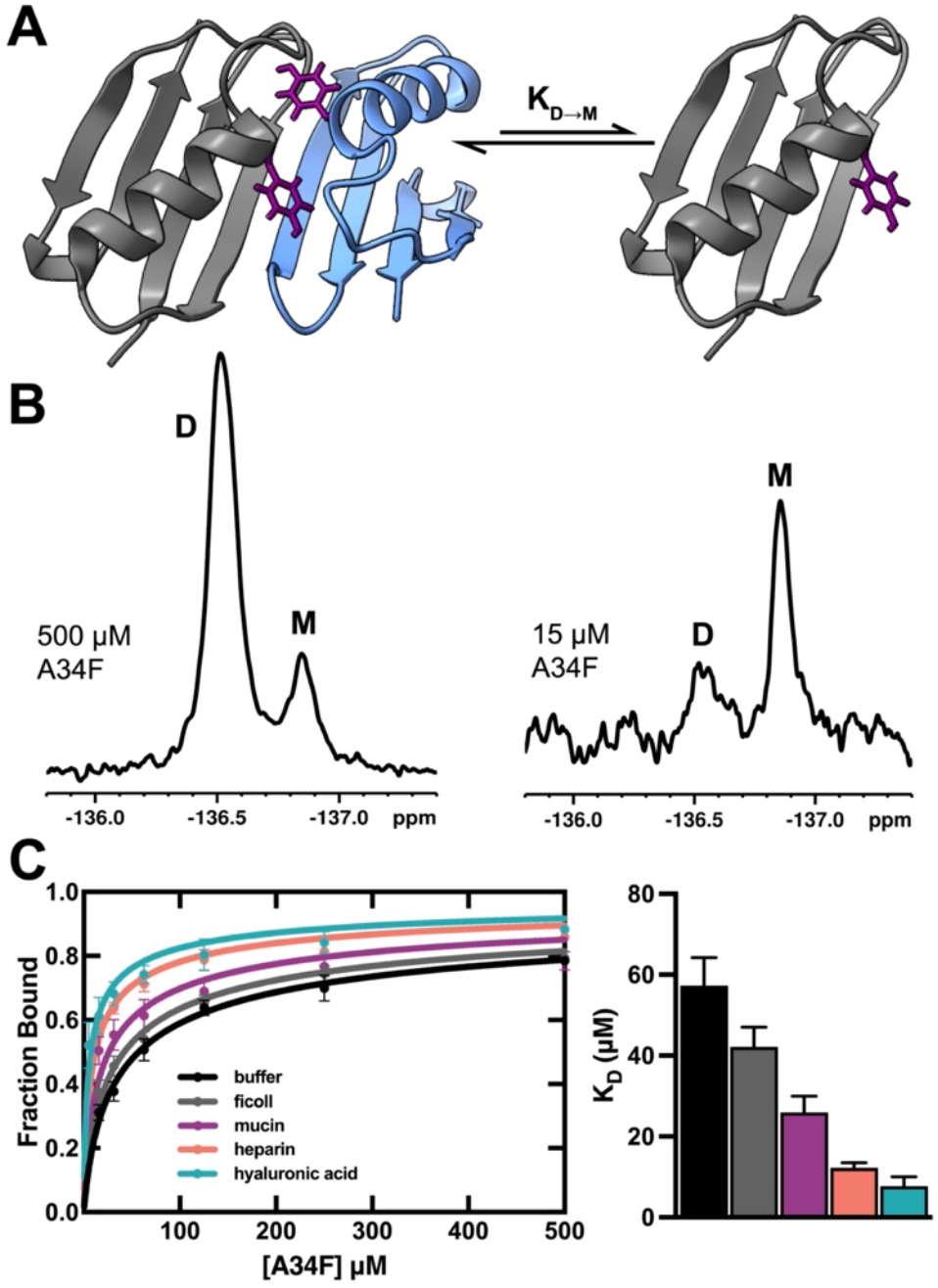
**a)** GB1 A34F (PDBID 2RMM) dimer and monomer. 3-fluoro-labelled Y33 residues are shown in purple. **b)** ^19^F NMR spectra of A34F with two Y33 resonances for the dimer (D) and monomer (M). Full spectra are shown in Fig. **S4**. Populations of each state change in a concentration-dependent manner. Spectra were taken in pH 7.0 buffer, 298K. **c)** binding curves from ^19^F NMR titrations of A34F in different crowding conditions (left) and comparison of dissociation constants in each condition (right). Data is shown as the mean ± SD of three measurements.

In buffer, we measured a K_D→M_ of 57 ± 6 µM, in excellent agreement with previous reports (Table **3** and Fig. **3c**).^15^ Stabilization of the dimer is minimal in ficoll (K_D→M_ = 42 ± 5 µM). The binding affinity of the dimer is enhanced by ~2 fold in 100 g/L mucin and ~4 fold in 100 g/L heparin. In hyaluronic acid, the K_D→M_ is 8 ± 2 µM and the dimer is stabilized by 1.2 ± 0.2 kcal/mol relative to buffer.

**Table 3.** Thermodynamic parameters for dimerization of A34F.

| | $K_{D \rightarrow M} (\mu\text{M})$ | $\Delta G_{D \rightarrow M}^{\circ'} (\text{kcal/mol})$ | $\Delta \Delta G_{D \rightarrow M}^{\circ'} (\text{kcal/mol})$ |
| --- | --- | --- | --- |
| **buffer** | $57 \pm 6$ | $4.4 \pm 0.1$ | - |
| **ficoll** | $42 \pm 5$ | $4.6 \pm 0.1$ | $0.2 \pm 0.1$ |
| **mucin** | $26 \pm 4$ | $4.9 \pm 0.1$ | $0.5 \pm 0.1$ |
| **heparin** | $12 \pm 1$ | $5.4 \pm 0.1$ | $1.0 \pm 0.1$ |
| **hyaluronic acid** | $8 \pm 2$ | $5.6 \pm 0.2$ | $1.2 \pm 0.2$ |

## DISCUSSION

Historically, discussions of macromolecular crowding were limited to hard-core repulsions, where proteins were treated as inert spheres inherently biased towards compacted structures in crowded environments.^2^ As protein stability was later measured in concentrated protein environments, cell lysates, and in living cells, classic crowding theories could not explain the enthalpic interactions between a test protein and its environment. Thus, modern crowding theory has evolved to understand that crowding includes an entropic packing component arising from volume exclusion, and an enthalpic chemical component arising from the electrostatic properties of a crowder and protein. Recent studies reflect the limited efficacy of synthetic sugar polymers as mimetics of the cellular interior, as these polymers lack charge functional groups.^5,18,44–46^ Inside cells, chemical interactions between protein surfaces have a greater influence on protein stability and interactions than excluded volume effects, and synthetic sugar polymers such as ficoll and dextran have been poor models of macromolecular crowding in the cytoplasm.^5^ Our work expands upon this paradigm, but using the component sugar polymers of the glycocalyx as crowders, which contain charged functional groups such as sulfates and carboxylic acids. We demonstrated how three physiological glycopolymers influence protein thermodynamics and kinetics, where chemical interactions were the dominant force. The uncharged polymer ficoll failed to mimic the crowding effect from charged physiologically glycopolymers, reflecting a need for more comprehensive models of macromolecular crowding which account for the chemical features of biological macromolecules.

SH3 and A34F are negatively charged proteins whose folding (SH3) and dimerization (A34F) are highly sensitive to charge-charge interactions. Both serve as useful models of how proteins may behave in the glycocalyx, with approximately half of human extracellular proteins weakly acidic (pI 4-7).^47^ We hypothesized that the negatively charged glycopolymers would enthalpically stabilize SH3 folding. By measuring the temperature dependence of the folding free energy, we were able to calculate the enthalpy 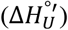 and entropy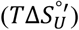for each condition, where the enthalpic stabilization outweighed the entropic destabilization. We observed an interaction enthalpy 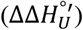 for SH3 folding of 4-6 kcal/mol for these physiologically relevant glycopolymers compared to the uncharged ficoll, whose interaction enthalpy was not distinguishable from dilute solution. Interestingly, we noted that hyaluronic acid was more stabilizing compared to SH3 than heparin. While both polymers have charged functional groups, hyaluronic acid is decorated with negatively charged carboxylate groups, while heparin is decorated with sulfates and was expected to have a greater negative charge and thus an expected greater effect. We observed that heparin had a greater 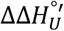 (5 ± 1) than the hyaluronic acid 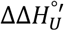(4 ± 1), but heparin also demonstrated a greater entropic destabilization than hyaluronic acid. As a result, hyaluronic acid demonstrated a larger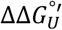 The differences of these effects may be explained by the size difference of the polymers, as the hyaluronic acid used in this study was 50 kDa and the heparin was 15 kDa. The interplay between polymer size, shape, and crowding effect is different between polymers and in different model systems, thus we are hesitant to overinterpret these data.^19,41,48^ We found that glycosylated mucins also stabilized SH3. The mucins used in this study were heterogenous as they were purified from porcine stomach.

Although they were less chemically defined than the GAGs, mucins are heavily sialylated and carry negative charge via the carboxylate functional groups present on sialic acid, similar to how GAGs convey negative surface charge through negatively charged chemical groups. These polymers generated stabilizing chemical interactions for SH3 as observed by the largest 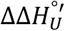 of the glycopolymers (6 ± 2), but mucin also had the largest entropic penalty to folding. These results demonstrate the potential for glycosylated proteins to generate stabilizing chemical environments for proteins and have effects comparable to negatively charged GAG polymers.

The mechanisms of SH3 stabilization by chemical interactions has been extensively studied in the context of protein crowders. *Smith* et al. demonstrated that crowders such as BSA and lysozyme has preferential interactions with the unfolded state of the protein as observed by changes in linewidths, a proxy for transverse relaxation rate T_2_, and by changes in the kinetic parameters of folding *k*_u_ and *k*_f_.^5^ Extending this analysis to our study, each glycopolymer we studied displayed a concentration dependent increase in linewidths for both the folded and unfolded states, however, the folded states show ~3x more broadening than the unfolded states, indictive of stabilizing interactions with the folded state (Table **2** and Figure **S2**). Using ^19^F-EXSY we measured the rates of folding and unfolding. We observed that hyaluronic acid and mucin increased the folding rate and decreased the unfolding rate, suggestive of interactions with the folded state (Table **2** and Figure **S3**). For heparin, these interactions are less clear; we saw a decrease in both the folding and unfolding rates, suggesting a balance between interactions with both states. Different from previously studied protein crowders that interact with the unfolded state of SH3, the biologically relevant glycopolymers we used stabilized SH3 folding via interactions with the folded state.

Extending these studies to protein-protein interactions, we used the model homodimer GB1 A34F to investigate how glycopolymer crowding affects protein dimerization. In physiological crowding environments, the A34F dimer was stabilized by 0.5-1.2 kcal/mol and binding of the dimer increased 2-6 fold. Following the same trends as protein folding, all three glycopolymers stabilized the dimer more than ficoll, and hyaluronic acid had the most significant effects. While we are unable to measure the interaction enthalpy of A34F, previous studies by Pielak and colleagues have demonstrated that A34F follows similar trends in chemical interactions to SH3, where charge-charge repulsions are stabilizing,^15,16,37^ suggesting that the stability of the dimer is also enthalpically motivated in this case. Importantly, this shows that the glycopolymers which compose the glycocalyx influence protein-protein interactions in the same fashion as protein folding.

The glycocalyx is a dynamic network of glycopolymers that is constantly changing as cells experience different stimuli.^21,49,50^ The data presented in this study demonstrate a potential mechanism in which the glycocalyx can regulate biological processes, as remodeling of the glycocalyx in response to stimuli or stress may alter the conformational landscapes of proteins native to the glycocalyx. Thus, if an extracellular domain of a signaling protein has a signaling inactive state A and a signaling active state B that undergo conformational change to drive a signaling process, under homeostatic conditions a tightly regulated thermodynamic barrier exists. As the glycocalyx gets remodeled, these barriers may shift, modulating the energetic differences between state A and B and either increasing or decreasing the thermodynamic barrier for these transitions. This presents an exciting avenue for further investigation.

## MATERIALS & METHODS

### Protein Expression & Purification

A pET-3a plasmid encoding the sequence for SH3 was transformed into BL21(DE3) *E. coli* cells and plated overnight on LB agar supplemented with 100 µg/mL carbenicillin. The next day, a single colony was used to inoculate 60 mL LB broth supplemented with the appropriate antibiotic. The culture was grown for 14 hours at 37°C. The next day, the overnight culture was split into the 3 × 1 L flasks of M9 minimal media supplemented with 100 µg/mL carbenicillin. Cells were grown at 37°C, 200 rpm to an OD of 0.6, at which point 60 mg 5-fluoroindole was added to each flask. Cultures continued to shake for 30 minutes, then were cooled to 18°C and protein expression was induced with isopropyl β-d-1-thiogalactopyranoside (IPTG) to a final concentration of 0.5 mM. Cultures were shaken at 18°C, 200 rpm for 16 hours, then centrifuged at 4000 x *g* for 20 minutes, and dry cell pellets were stored at –80°C.

Protein purification was performed as previously described.^51^ Cell pellets were resuspended in lysis buffer (20 mM TRIS pH 8.5), lysed by sonication, and clarified by centrifugation at 24,000 x *g* for 30 minutes. The soluble material was loaded on a HiTrap Q HP column on an ÄKTA go™ system, washed in lysis buffer, and eluted in the same buffer with a gradient of 0 – 1 M NaCl. Fractions containing SH3 were pooled, concentrated to 1 mL, and further purified by size exclusion chromatography on a Hiload Superdex 75 16/600 column. Pure protein fractions were dialyzed into 18.2 MΩ cm^−1^ water for 12 hours and concentrated to < 2 mLs at 4°C, 4000 x *g* in a 5 kDa cutoff concentrator. Final protein purity was determined by SDS-PAGE and ESI-MS (expected mass 6878 Da, observed 6879 Da), and concentration was determined by nanodrop (8480 M^−1^cm^−1^) (Fig. **S5**). Samples were aliquoted for NMR, lyophilized, and stored at –80°C.

A pET-3a plasmid containing the sequence for the T2Q A34F variant of GB1 (here referred to as A34F) was transformed into BL21(DE3) *E. coli* cells. Expression of GB1 followed the procedure described above with two exceptions. A34F was grown in ^15^N enriched M9 media, and 60 mg 3-fluorotyrosine was added to cell cultures at an OD of 0.6 instead of 5-fluorindole. A34F was purified as previously described^15^ following the same procedure described in the paragraph above. Protein concentration was determined by nanodrop (9970 M^−1^cm^−1^) and purity was confirmed by SDS-PAGE and ESI-MS (expected mass 6393 Da, observed 6417 Da) (Fig. **S6**).

### NMR

Fluorine labeled SH3 was resuspended in 20 mM phosphate pH 7.0 in 10% D_2_O to a final concentration of 200 µM with the appropriate glycopolymer. Heparin sodium (15 kDa), ficoll (70 kDa), and mucin were purchased from Sigma. Hyaluronic acid (50 kDa) was purchased from Fisher.

Experiments were performed on a Bruker Ascend Evo spectrometer operating at a ^19^F Larmor frequency of 565 MHz equipped with a cryogenic TCI probe tunable to fluorine. 1D spectra of SH3 were acquired with an acquisition of 4096 points and a relaxation delay of 2 s, with an offset of −120 ppm and a sweep width of 30 ppm. The number of scans was either 128 or 256. For variable temperature experiments, spectra were collected at 288K, 293K, 298K, 303K, 308K, 313K, and 318K. Spectra were acquired at 298K before and after the temperature gradient to ensure reversibility of folding. For folding experiments, SH3 was suspended in the same buffer to a final concentration of 400 µM. A Bruker library NOESY experiment (t_mix_ = 20 ms, 75 ms (x3), 125 ms, 175 ms, 250 ms, 400 ms, 600 ms, and 800 ms) was used to measure folding and unfolding rates. 2048 points were acquired in the f2 dimension with 128 points in the f1 dimension, across a sweep width of 10 ppm. 16 transients were acquired per increment, and the relaxation delay was 2 s. Titrations with GB1 were done using the same instrument. Protein was resuspended in 20 mM phosphate, pH 7.0 in 10% D_2_O to a concentration of 500 µM and serial diluted to 250 µM, 125 µM, 62.5 µM, 31.2 µM, and 15.6 µM in the same condition. 1D ^19^F spectra were acquired with an acquisition of 4096 points and a relaxation delay of 2 s, with an offset of −130 ppm and a sweepwidth of 30 ppm. The number of scans varied from 128 to 2048, depending on the protein concentration and crowder conditions.

Data were processed on Topspin 4.5.0. An exponential line broadening function of 10 Hz was applied to each FID before Fourier transformation. For 1D experiments with SH3, peak integrations were used to determine ρ_U_ and ρ_F_. For exchange experiments, the ratio of peak volumes was extracted, and the rates of exchange were calculated as described by Gustafson et al.^43^ One mixing time was measured three times and the sample standard deviation was used to drive a Monte Carlo analysis (*n* = 1000) to propagate error to the rest of the data set.^5^ For 1D experiments with GB1, Y33 peak integrations were used to determine *f*_M_ and *f*_D_. Additional details of data fitting are described in the results section.

## ACKNOWLEDGEMENTS

This work was supported by R00GM145970 to A.J.G. and U54CA272220. We thank Gary Pielak and Annelise Gorensek-Benitez for useful discussion. We thank Xuemei Huang and Anthony Mrse for maintaining the spectrometers.

## Author contributions

**Alex J. Guseman:** conceptualization; investigation; writing-review and editing; funding acquisition, formal analysis; writing-original draft. **Sabrina M. Richter:** conceptualization; investigation; writing-review and editing; formal analysis; writing-original draft. **Neal Brook: i**nvestigation; writing-review and editing.

