## Supplementary material for "Glycopolymers stabilize protein folding and protein-protein interactions via enthalpic interactions": Figure S1

**^1^Department of Chemistry and Biochemistry University of California San Diego, La Jolla California**

***Corresponding Author**

**Table S1.** Changes in thermodynamic stability of SH3 in crowding environments.

|  | $\boldsymbol{\Delta G}_{\boldsymbol{U}}^{\boldsymbol{^{\circ}'}}$ **(kcal/mol)** | $\boldsymbol{\Delta\Delta G}_{\boldsymbol{U}}^{\boldsymbol{^{\circ}'}}$ **(kcal/mol)** |
| --- | --- | --- |
| **buffer** | 0.50 ± 0.06 | - |
| **ficoll (g/L)** |  |  |
| 50 | 0.54 ± 0.01 | 0.04 ± 0.06 |
| 100 | 0.60 ± 0.02 | 0.11 ± 0.06 |
| 200 | 0.69 ± 0.05 | 0.20 ± 0.07 |
| 300 | 0.81 ± 0.06 | 0.31 ± 0.08 |
| **heparin (g/L)** |  |  |
| 50 | 0.61 ± 0.03 | 0.12 ± 0.06 |
| 100 | 0.82 ± 0.01 | 0.33 ± 0.06 |
| 200 | 1.08 ± 0.04 | 0.58 ± 0.07 |
| 300 | 1.41 ± 0.06 | 0.92 ± 0.08 |
| **hyaluronic acid (g/L)** |  |  |
| 50 | 0.86 ± 0.05 | 0.36 ± 0.08 |
| 100 | 0.99 ± 0.01 | 0.49 ± 0.06 |
| 200 | 1.6 ± 0.1 | 1.1 ± 0.2 |
| 300 | 2.0 ± 0.1 | 1.5 ± 0.1 |
| **mucin (g/L)** |  |  |
| 50 | 0.70 ± 0.01 | 0.21 ± 0.06 |
| 100 | 0.85 ± 0.01 | 0.36 ± 0.06 |
| 200 | 1.13 ± 0.06 | 0.64 ± 0.08 |

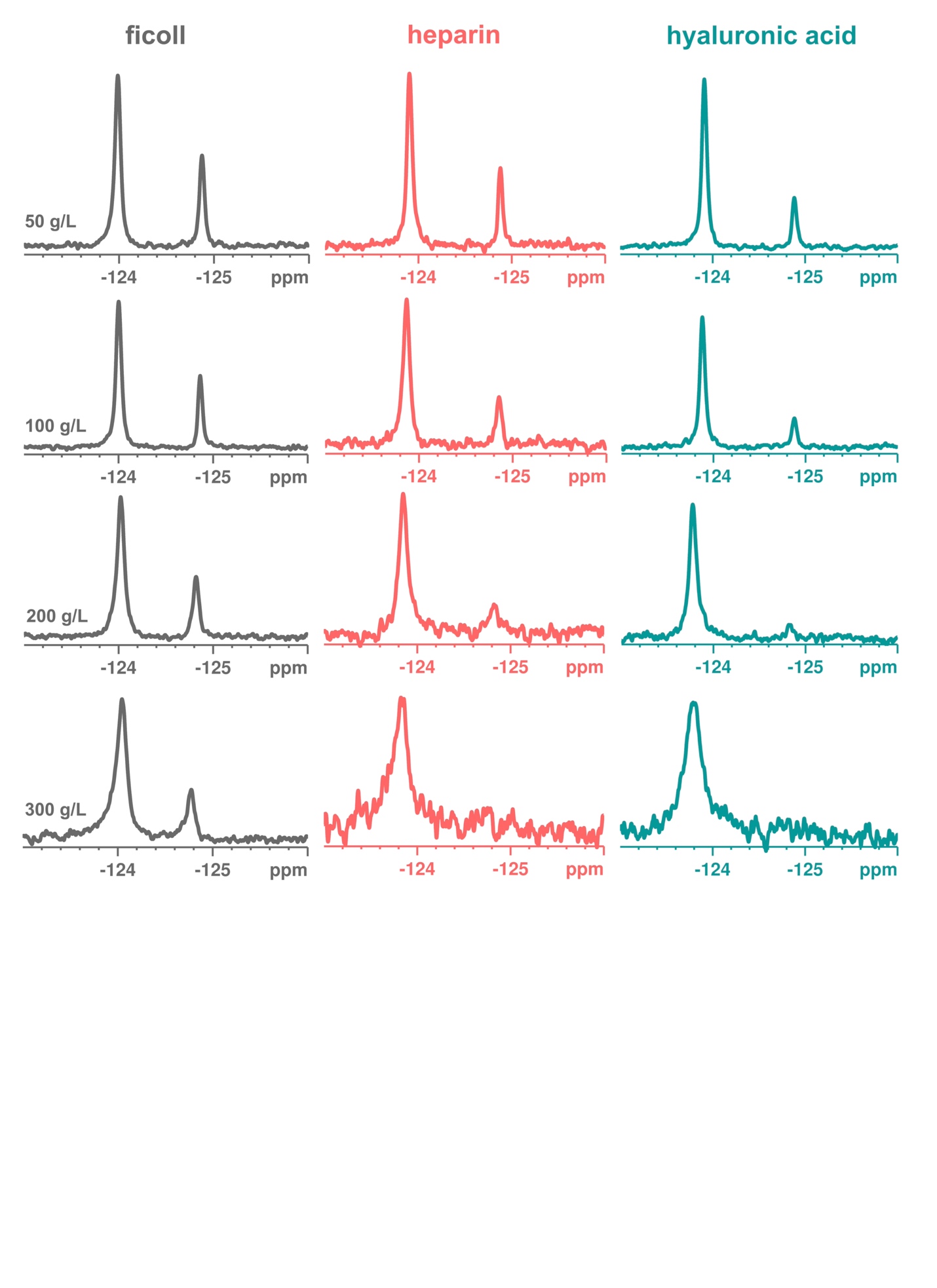
**Figure S1.** Representative ^19^F NMR spectra of 5-fluorotryptophan-labelled SH3 in ficoll, heparin, and hyaluronic acid. The upfield resonance corresponds to the unfolded population and the downfield resonance to the folded population. Assignments are as previously published.^1^ All spectra were collected at pH 7.0, 298K.

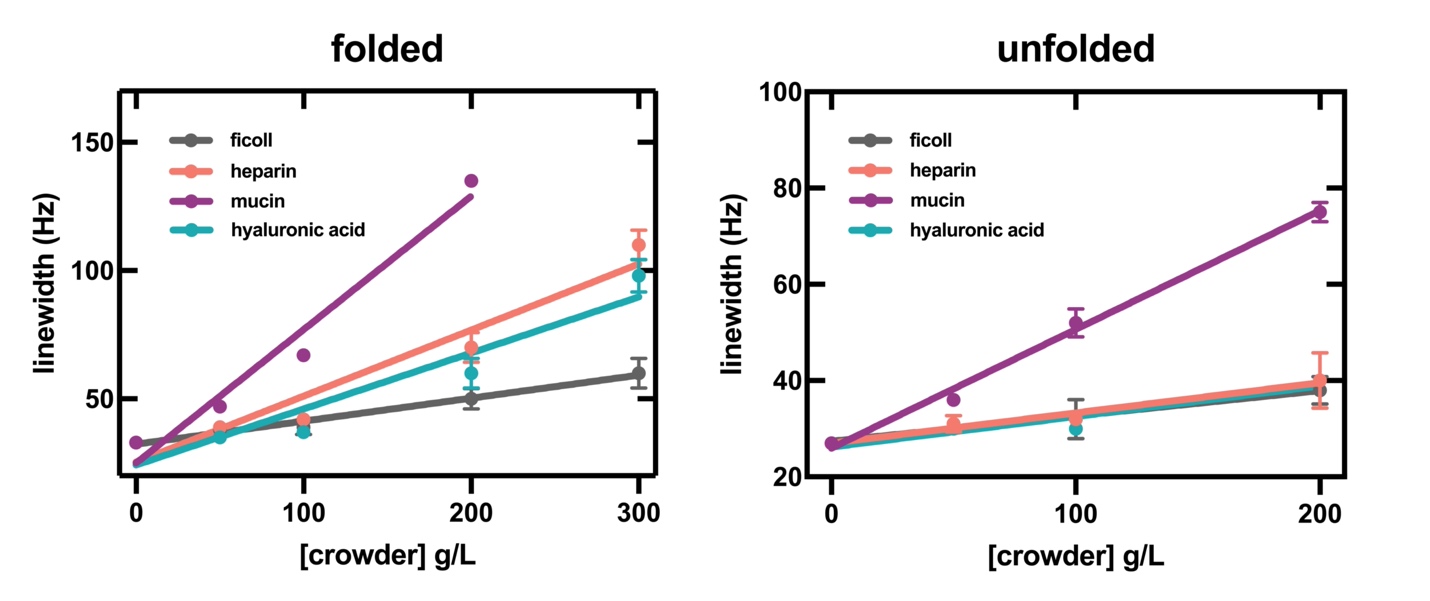

**Figure S2.** Linewidths of folded and unfolded state peaks in each condition at increasing crowder concentrations. Unfolded peak linewidths could not be determined above 200 g/L because the folded state is strongly biased. Data are shown as the mean ± SD of three measurements.

**Table 2.** Linewidths of SH3 folded and unfolded resonances and estimated T2 values^2^ in each condition.

|  |  | **linewidth (Hz)** | |  | **T2 (ms)** | |
| --- | --- | --- | --- | --- | --- | --- |
|  |  | **folded** | **unfolded** |  | **folded** | **unfolded** |
| **buffer** |  | 33 ± 1 | 27 ± 1 |  | 9.6 ± 0.3 | 11.8 ± 0.4 |
| **ficoll (g/L)** |  |  |  |  |  |  |
| 50 |  | 38 ± 1 | 31 ± 1 |  | 8.4 ± 0.2 | 10.3 ± 0.3 |
| 100 |  | 39 ± 5 | 32 ± 7 |  | 8 ± 1 | 10 ± 2 |
| 200 |  | 50 ± 7 | 38 ± 5 |  | 6.4 ± 0.9 | 8 ± 1 |
| 300 |  | 60 ± 10 | 40 ± 6 |  | 5.3 ± 0.9 | 8 ± 1 |
| **heparin (g/L)** |  |  |  |  |  |  |
| 50 |  | 39 ± 2 | 31 ± 3 |  | 8.2 ± 0.4 | 10 ± 1 |
| 100 |  | 42 ± 2 | 32 ± 1 |  | 7.6 ± 0.4 | 10.0 ± 0.3 |
| 200 |  | 70 ± 10 | 40 ± 10 |  | 4.6 ± 0.6 | 8 ± 2 |
| 300 |  | 110 ± 10 | n.d. |  | 2.9 ± 0.3 | n.d. |
| **hyaluronic acid (g/L)** |  |  |  |  |  |  |
| 50 |  | 35 ± 1 | 28 ± 1 |  | 9.1 ± 0.3 | 11.4 ± 0.4 |
| 100 |  | 37 ± 1 | 30 ± 2 |  | 8.6 ± 0.2 | 10.6 ± 0.7 |
| 200 |  | 60 ± 10 | 40 ± 10 |  | 5.3 ± 0.9 | 8 ± 2 |
| 300 |  | 98 ± 9 | NA |  | 3.2 ± 0.3 | n.d. |
| **mucin (g/L)** |  |  |  |  |  |  |
| 50 |  | 47 ± 1 | 36 ± 2 |  | 6.8 ± 0.1 | 8.8 ± 0.5 |
| 100 |  | 67 ± 1 | 52 ± 5 |  | 4.8 ± 0.1 | 6.1 ± 0.6 |
| 200 |  | 135 ± 3 | 100 ± 30 |  | 2.4 ± 0.1 | 3 ± 1 |

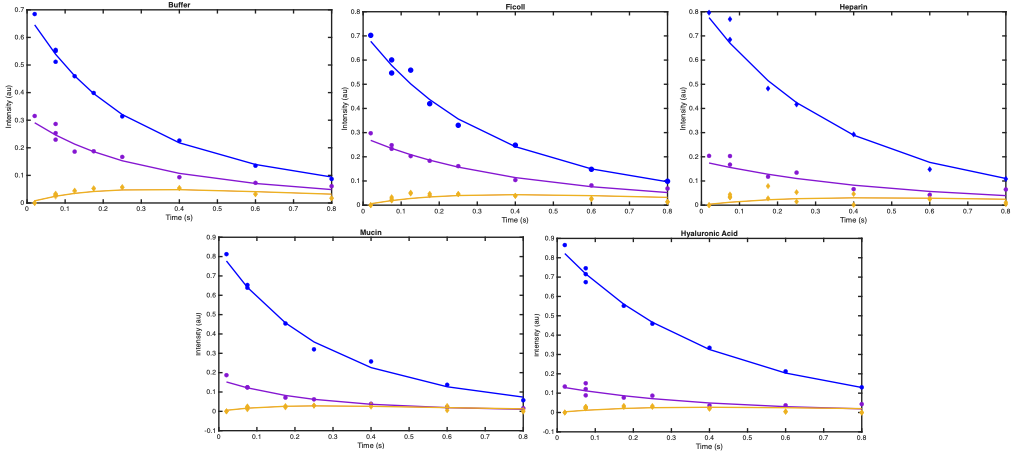

**Figure S3**. Representative EXSY curves for SH3 folding and unfolding in each glycopolymer at 100 g/L and fit to the EXSY model established by Farrow et al.^3^ Each curve represents the change in intensity over time for the fold state (blue), unfolded state (purple), and cross peaks (gold).

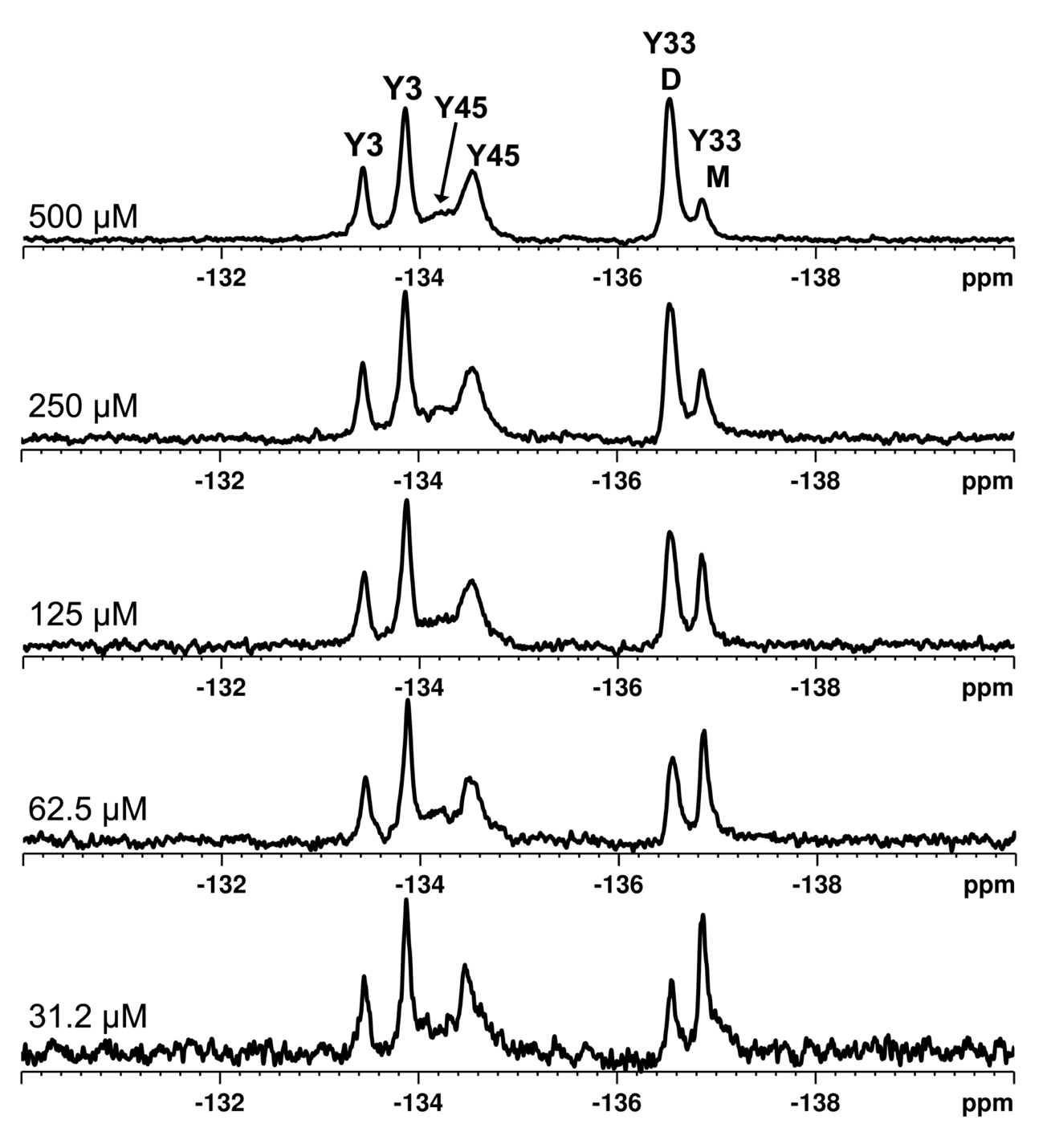

**Figure S4.** Representative ^19^F NMR titration of GB1. Spectra of 3-fluorotyrosine labelled GB1 show six peaks in total, two for each tyrosine. The rotamers of Y3 and Y45 are in slow exchange, and each show two resonances. Y33 shows two resonances, and peak ratios change with protein concentration. The upfield resonance corresponds to the monomer (M) and the downfield resonance to the dimer (D). Assignments are as previously published.^4^ All spectra were taken at pH 7.0, 298K.

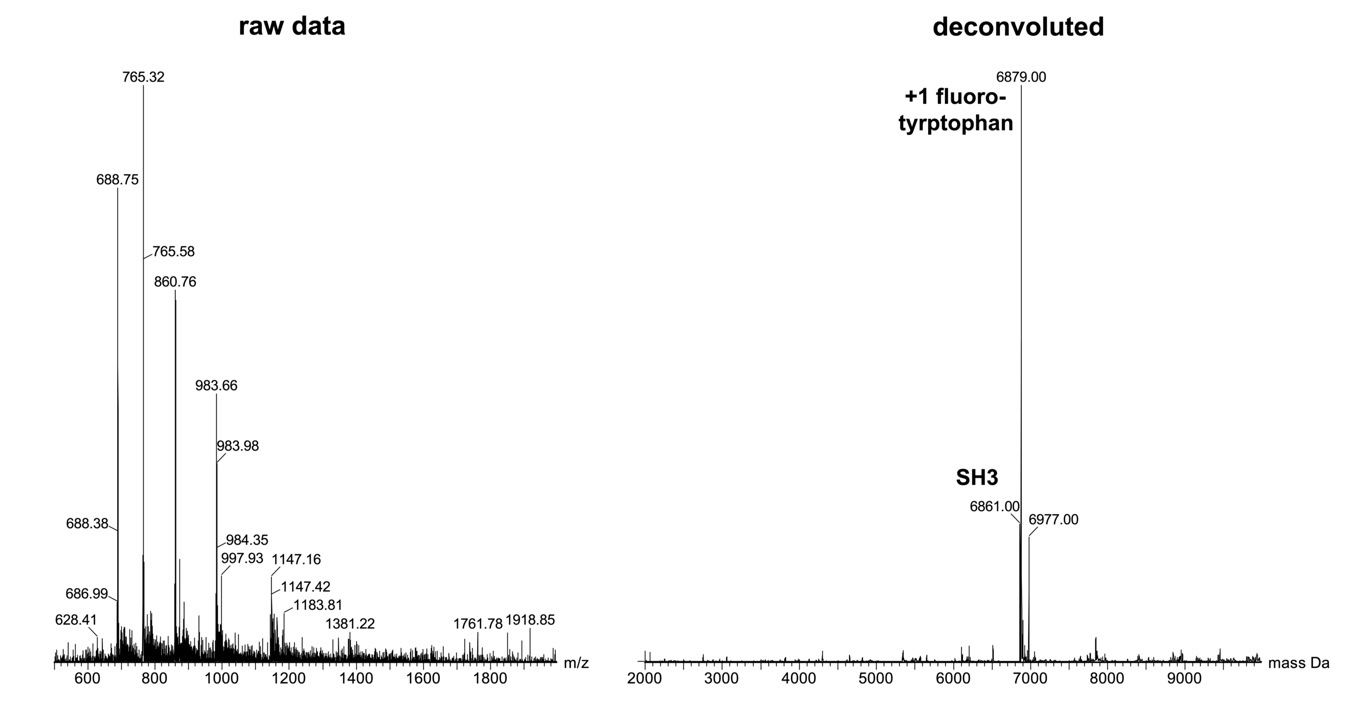

**Figure S5.** Raw (left) and deconvoluted (right) ESI-MS of SH3 labelled with 5-fluoro-tyrptophan. Peaks indicate protein with 0 (6861 Da) and 1 (6879 Da) fluorinated residue.

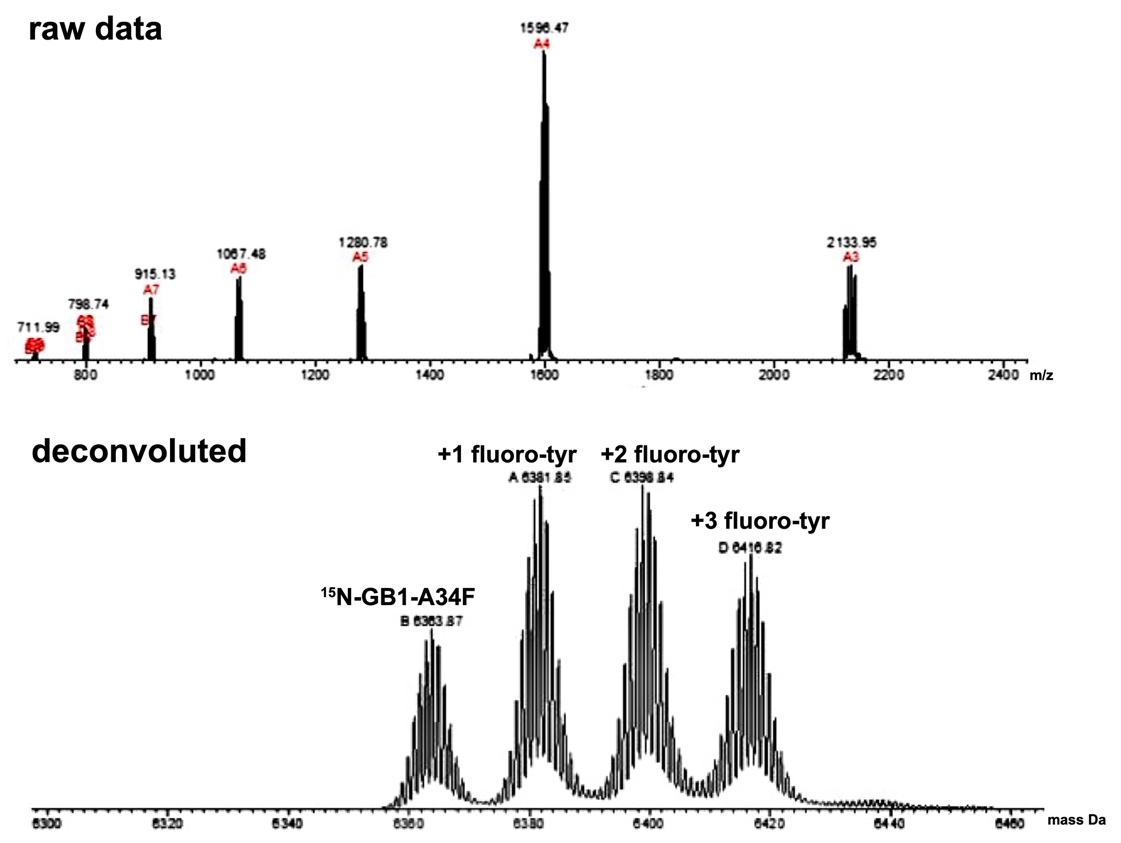

**Figure S6.** Raw (top) and deconvoluted (bottom) ESI-MS of ^15^N-enriched GB1 A34F labelled with 3-fluorotyrosine. Peaks indicate protein with 0 (6364 Da), 1 (6382 Da), 2 (6399 Da), and 3 (6417 Da) fluorinated tyrosine residues.

(1) Evanics, F.; Bezsonova, I.; Marsh, J.; Kitevski, J. L.; Forman-Kay, J. D.; Prosser, R. S. Tryptophan Solvent Exposure in Folded and Unfolded States of an SH3 Domain by^19^ F and^1^ H NMR. *Biochemistry* **2006**, *45* (47), 14120–14128.
(2) Mladenov, G.; Dimitrov, V. S. Extraction of *T*_2_ from NMR Linewidths in Simple Spin Systems by Use of Reference Deconvolution. *Magnetic Reson in Chemistry* **2001**, *39* (11), 672–680.
(3) Farrow, N. A.; Zhang, O.; Forman-Kay, J. D.; Kay, L. E. A Heteronuclear Correlation Experiment for Simultaneous Determination of 15N Longitudinal Decay and Chemical Exchange Rates of Systems in Slow Equilibrium. *J Biomol NMR* **1994**, *4* (5), 727–734.
(4) Ye, Y.; Liu, X.; Zhang, Z.; Wu, Q.; Jiang, B.; Jiang, L.; Zhang, X.; Liu, M.; Pielak, G. J.; Li, C. ^19^ F NMR Spectroscopy as a Probe of Cytoplasmic Viscosity and Weak Protein Interactions in Living Cells. *Chemistry A European J* **2013**, *19* (38), 12705–12710.
